# Humanized *Drosophila* model of the Meier-Gorlin syndrome reveals conserved and divergent features of the Orc6 protein

**DOI:** 10.1101/2020.04.29.068866

**Authors:** Maxim Balasov, Katarina Akhmetova, Igor Chesnokov

## Abstract

Meier-Gorlin syndrome (MGS) is a rare autosomal recessive disorder characterized by microtia, primordial dwarfism, small ears and skeletal abnormalities. Patients with MGS often carry mutations in the genes encoding the subunits of the Origin Recognition Complex (ORC), components of the pre-replicative complex (pre-RC) and replication machinery. Orc6 is an important component of ORC and has functions in both DNA replication and cytokinesis. A mutation in conserved C-terminal motif of Orc6 associated with MGS impedes the interaction of Orc6 with core ORC. Recently, new mutation in Orc6 was also identified however, it is localized in the N-terminal domain of the protein. In order to study the functions of Orc6 we used human gene to rescue the *orc6* deletion in *Drosophila*. Using the “humanized” Orc6-based *Drosophila* model of the Meier-Gorlin syndrome we discovered that unlike previous Y225S MGS mutation in Orc6, the K23E substitution in the N-terminal TFIIB-like domain of Orc6 disrupts the protein ability to bind DNA. Our studies revealed the importance of evolutionary conserved and variable domains of Orc6 protein and allowed the studies of human protein functions and the analysis of the critical amino acids in live animal heterologous system as well as provided novel insights into the mechanisms underlying MGS pathology.

## Introduction

DNA replication is fundamentally important for tissue development and homeostasis and impairments of the DNA replication machinery can have catastrophic consequences for genome stability and cell division. Meier-Gorlin Syndrome (MGS) is an autosomal recessive disorder that is also known as ear, patella, short stature syndrome and/or microtia, absent patella, micrognathia syndrome, highlighting the core clinical phenotypes (Gorlin et al. 1975), (de Munnik et al. 2012b), (de Munnik et al. 2012a), (Bicknell et al. 2011a). The genes affected by MGS mutations include many members of pre-replicative complex (pre-RC), such as ORC subunits (Orc1, Orc4, Orc6), Cdc6, Cdt1, CDC45, MCM5 as well as Geminin (Bicknell et al. 2011a), (Bicknell et al. 2011b), (Guernsey et al. 2011), (Fenwick et al. 2016), (Vetro et al. 2017), (Burrage et al. 2015), (McDaniel et al. 2020) suggesting that the clinical phenotype is caused by defects in DNA replication initiation. As pre-RC complex is essential for DNA replication, the mutations in its components are expected to impair cell proliferation and to reduce growth.

The Origin Recognition Complex (ORC) plays a central role in the initiation of DNA replication but is also involved in non-replicative functions (Bell and Stillman 1992), (Bell 2002), (Sasaki and Gilbert 2007), (Chesnokov 2007). The smallest subunit of ORC, Orc6 is the most divergent and possibly the most enigmatic among ORC subunits. Orc6 is important for DNA replication in all species (Lee and Bell 1997), (Chesnokov et al. 2001), (Chesnokov et al. 2003), (Semple et al. 2006), (Chen et al. 2007), (Balasov et al. 2007), (Balasov et al. 2009). It is also essential for cytokinesis in *Drosophila* and human cells (Prasanth et al. 2002), (Chesnokov et al. 2003), (Bernal and Venkitaraman 2011). In *Drosophila*, Orc6 stimulates septin complex GTPase activity and polymerization during filament assembly through protein-protein interactions (Huijbregts et al. 2009), (Akhmetova et al. 2015). Metazoan Orc6 proteins consist of two functional domains: a larger N-terminal domain important for binding of DNA and smaller C-terminal domain important for protein-protein interactions (Chesnokov et al. 2003), (Balasov et al. 2007), (Duncker et al. 2009), (Bleichert et al. 2013). It has been shown that in metazoan species N-terminal domain of Orc6 carries a structural homology with TFIIB transcription factor (Chesnokov et al. 2003), (Balasov et al. 2007), (Liu et al. 2011). The conserved motif in C-terminus of Orc6 is responsible for the interaction with Orc3 subunit of ORC (Bleichert et al. 2013) and tyrosine 232 to serine mutation in this region of the protein is linked to the MGS in humans (Bicknell et al. 2011a), (de Munnik et al. 2012a). In our earlier studies we introduced MGS mutation in *Drosophila* Orc6 (Y225S) and established fruit fly model of MGS. Obtained flies displayed severe replication and developmental defects due to impaired interaction of mutant Orc6 protein with the rest of the ORC (Balasov et al. 2015).

Recently, new Orc6 mutation was described (Li et al. 2017) that also resulted in MGS. Unlike previously described MGS mutation (Bicknell et al. 2011a), this amino acid substitution (Lys23Glu) localizes in the N-terminal domain of Orc6 which is important for DNA binding. In our study, in order to investigate the functions of different Orc6 domains in live animal model we introduced human, *Drosophila*, as well as hybrid *human-Drosophila orc6* genes into flies with *orc6* null mutant background. Using “humanized” fly strains carrying K23E MGS mutation, we identified a molecular mechanism resulting in MGS phenotype and compared them with our early MGS fly model carrying Y225S in the C-terminus of Orc6. We found that despite different underlying molecular mechanisms both MGS mutations result in similar phenotypes, deficient pre-RC formation and reduced DNA replication.

## RESULTS

### 1. Establishing *Drosophila* strains to study the functions of human Orc6 *in vivo*

In our previous work we generated a deletion of the *orc6* gene in *Drosophila* allowing the studies of the protein *in vivo*. We showed that the *orc6* deletion resulted in defects in DNA replication as well as abnormal chromosome condensation and segregation (Balasov et al. 2009). Orc6 is the least conserved among ORC subunits, however, metazoan Orc6 proteins show significantly higher homology as compared to budding yeast protein. We asked if an expression of human Orc6 protein may rescue third instar lethality associated with a deletion of *orc6* gene in *Drosophila*. Full length human *orc6* gene under control of native *orc6* promoter was not able to rescue a lethality of the Orc6 deficient flies (**Figure 1A** and (Balasov et al. 2009)). To investigate the functions of the protein in more detail, additional constructs were created (**Figure 1A**). The metazoan Orc6 consists of two domains, a larger N-terminal domain which carries homology with TFIIB transcription factor and is important for DNA binding (Chesnokov et al. 2003), (Balasov et al. 2007), (Liu et al. 2011) and a shorter C-terminal domain important for the function of Orc6 in cytokinesis (Prasanth et al. 2002), (Chesnokov et al. 2003), (Huijbregts et al. 2009), (Bernal and Venkitaraman 2011), (Akhmetova et al. 2015). The C-terminal domain of metazoan Orc6 is also essential and sufficient for the interaction of the protein with the core ORC through its Orc3 subunit (Bleichert et al. 2013). Two hybrid constructs were created: the first consisted of *Drosophila* N-terminal TFIIB like domain and human C-terminus (Orc6-DH); the second construct contained human N-terminal domain and *Drosophila* C-terminus (Orc6-HD). Again, we tested the constructs for their ability to rescue *orc6* deletion. The expression of Orc6-DH did not rescue the lethality, however, the expression of Orc6-HD resulted in a full rescue and obtained flies were undistinguishable from the flies rescued with wild type *Drosophila* Orc6 (**Figure 1A**).

**Figure 1.**
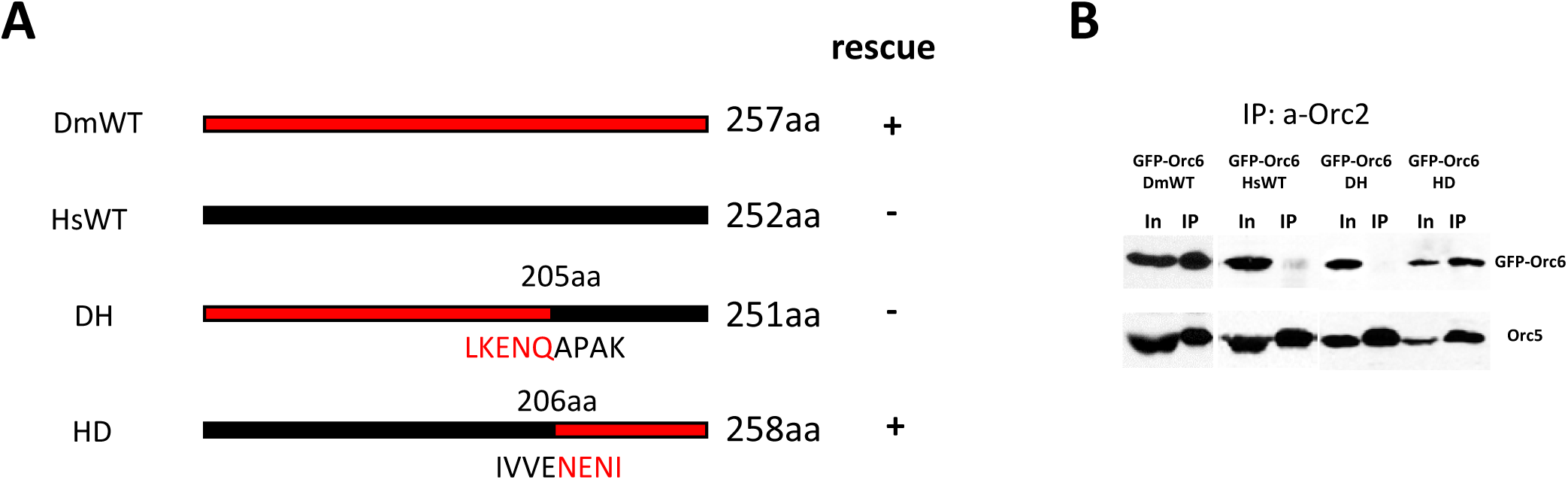
Design of hybrid Orc6s. **(A)** Cartoon diagrams of transgenic constructs. Black color represents Human Orc6, red color corresponds to *Drosophila* Orc6. DmWT and HsWT – wild type *Drosophila* and human proteins, respectively. DH – hybrid protein containing first 205 amino acids of *Drosophila* and last 46 amino acids of human Orc6. HD – hybrid protein containing first 205 amino acids of human and last 52 amino acids of *Drosophila* Orc6. **(B)** Immunoprecipitation of the ORC complex with anti-Orc2 antibodies from ovaries expressing GFP-tagged Orc6s shown in (A).

Next, we analyzed the ability of human and hybrid Orc6 proteins to associate with ORC complex in heterozygous flies. We found that Orc6-HD but not Orc6-DH could be immunoprecipitated with anti-Orc2 antibodies from ovary extracts (**Figure 1B**). Small amounts of human Orc6 protein could be detected in the IP material from strains carrying GFP-Orc6-Hs possibly reflecting loose association of human Orc6 with the core ORC (**Figure 1B**). Consistent with these results, DNA replication judged by BrdU incorporation was reduced in larval brains of the flies carrying Orc6-Hs or Orc6-DH constructs (**Figure 2 B**, **D**) but was similar to the wild type Orc6-Dm in the case of Orc6-HD hybrid construct (**Figure 2 A**, **C**). Furthermore, the MCM helicase association with the chromatin was diminished in Orc6-DH but not in Orc6-HD strains (**Supplementary Figure 1A**).

**Figure 2.**
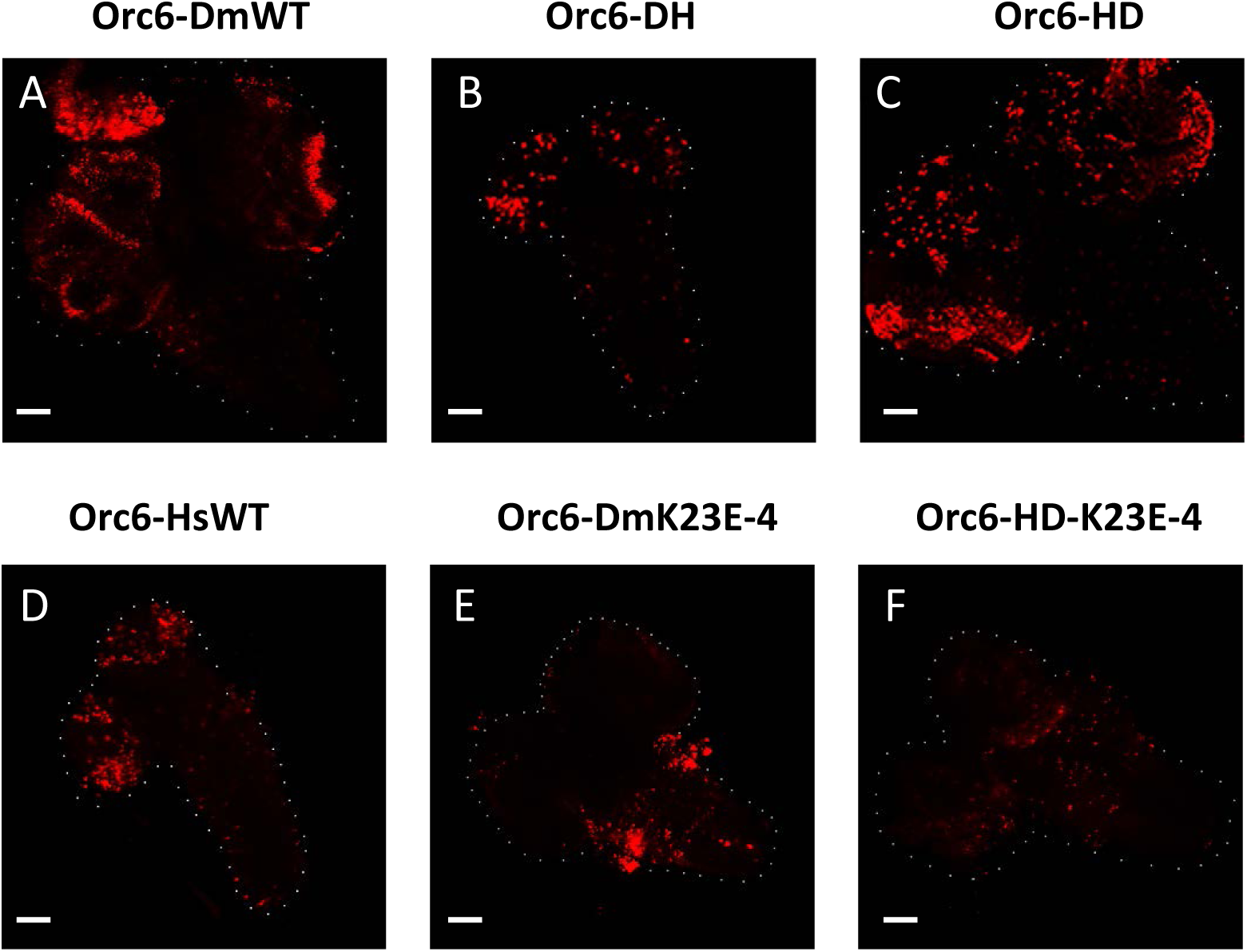
BrdU incorporation in larval brains. Third instar larval brains of *Drosophila orc6* deletion rescued with different *orc6* transgenes are shown. Transgenes are as follows: **(A)** Wild type *Drosophila* Orc6. **(B)** Orc6-DH hybrid. **(C)** Orc6-HD hybrid. **(D)** Wild type human Orc6. **(E)** *Drosophila* Orc6-K23E mutant. **(F)** Orc6-HD hybrid with K23E mutation. Scale bar represents 50 µm.

We also analyzed mitotic chromosomes from neural ganglia derived from larvae carrying mutant and wild type *orc6* genes. Normal *Drosophila melanogaster* karyotype consists of four pairs of chromosomes (**Figure 3A**, **Orc6-Dm wild type**). In larvae carrying *orc6* deletion, or rescued with Orc6-DH (**Figure 3A**) or Orc6-HsWT (**Figure 3C**) constructs, chromosomes lost parts of their arms, appeared aberrantly condensed, fragmented, misaligned and sometimes polyploid. Orc6-HD transgene expression restored both DNA replication and mitotic chromosome structure (**Figure 2C and 3A**). Since Orc6-HD compensated *orc6* deletion similar to the wild type *Drosophila* Orc6, this transgene can be used for studying mutations in the human large N-terminal TFIIB like domain portion of Orc6 protein in live animal model.

**Figure 3.**
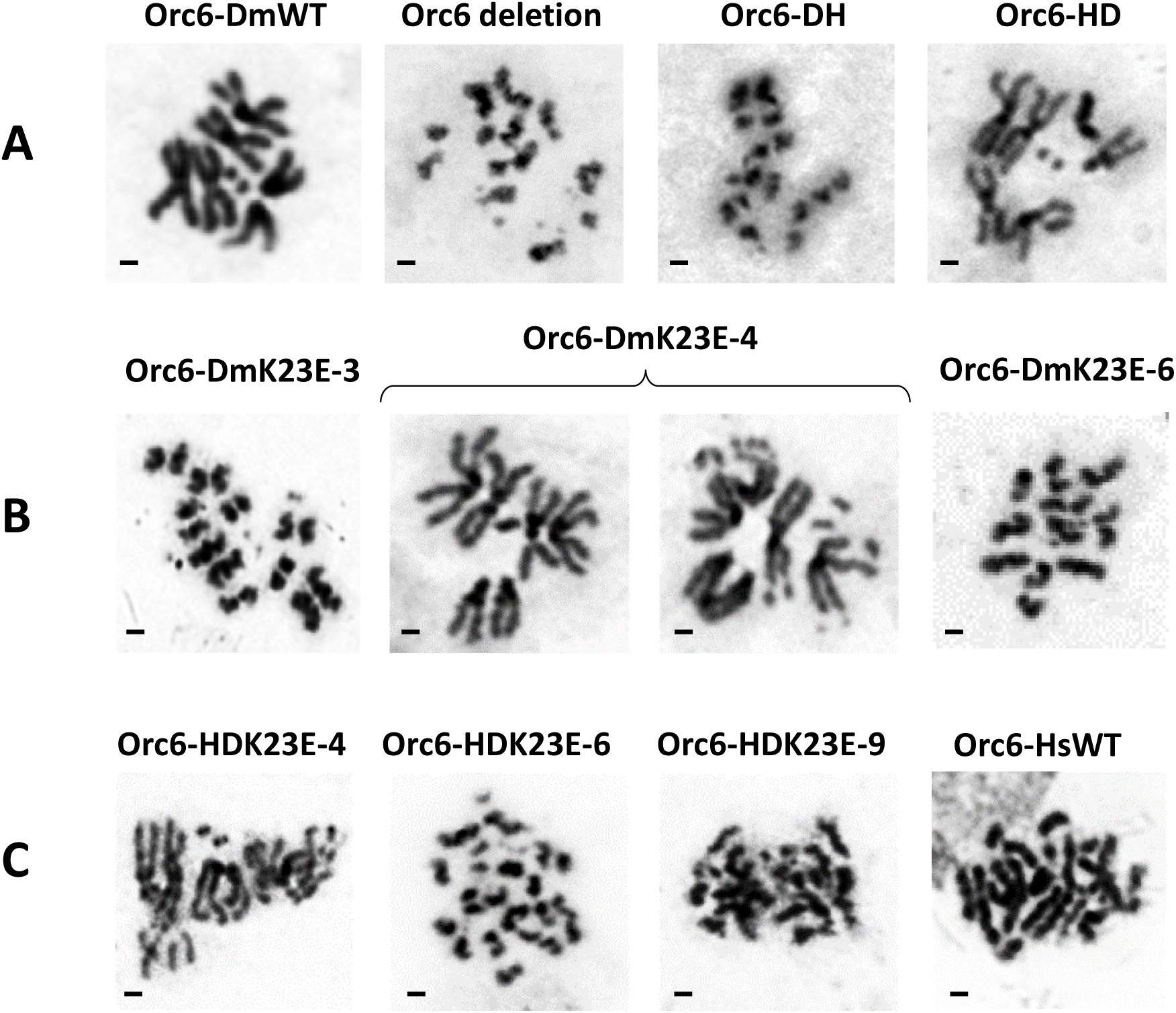
Mitoses in larval neuroblasts. *Orc6* deletion flies were rescued with specified transgenes and third instar larvae brains were analyzed for the karyotype. **(A)** Orc6 deletion mutant and hybrid Orc6s (Orc6-DH, Orc6-HD) compared to *Drosophila* wild type (Orc6-DmWT). **(B)** Three independent transgenes bearing K23E mutation (Orc6-DmK23E-3, Orc6-DmK23E-4, Orc6-DmK23E-6). **(C)** Three independent hybrid Orc6-HD transgenes bearing mutation in human portion of Orc6 (Orc6-HDK23E-4, Orc6-HDK23E-6, Orc6-HDK23E-9) compared to human wild type Orc6 (Orc6-HsWT). Scale bar represents 1 µm.

### 2. The analysis of flies carrying MGS mutation localized in the N-terminal domain of Orc6

Recently, the first diagnosed MGS case in China was described (Li et al. 2017). The patient has featured a short stature, microtia, small patella and craniofacial abnormalities characteristic for the Meier-Gorlin syndrome. It was reported that the patient carried a novel homozygous mutation in the *orc6* gene resulting in a change of A to G at position 67 which corresponds to the conversion of Lys at position 23 to Glu. Interestingly, this mutation is localized within N-terminal TFIIB like domain of the protein (**Figure 4A**).

**Figure 4.**
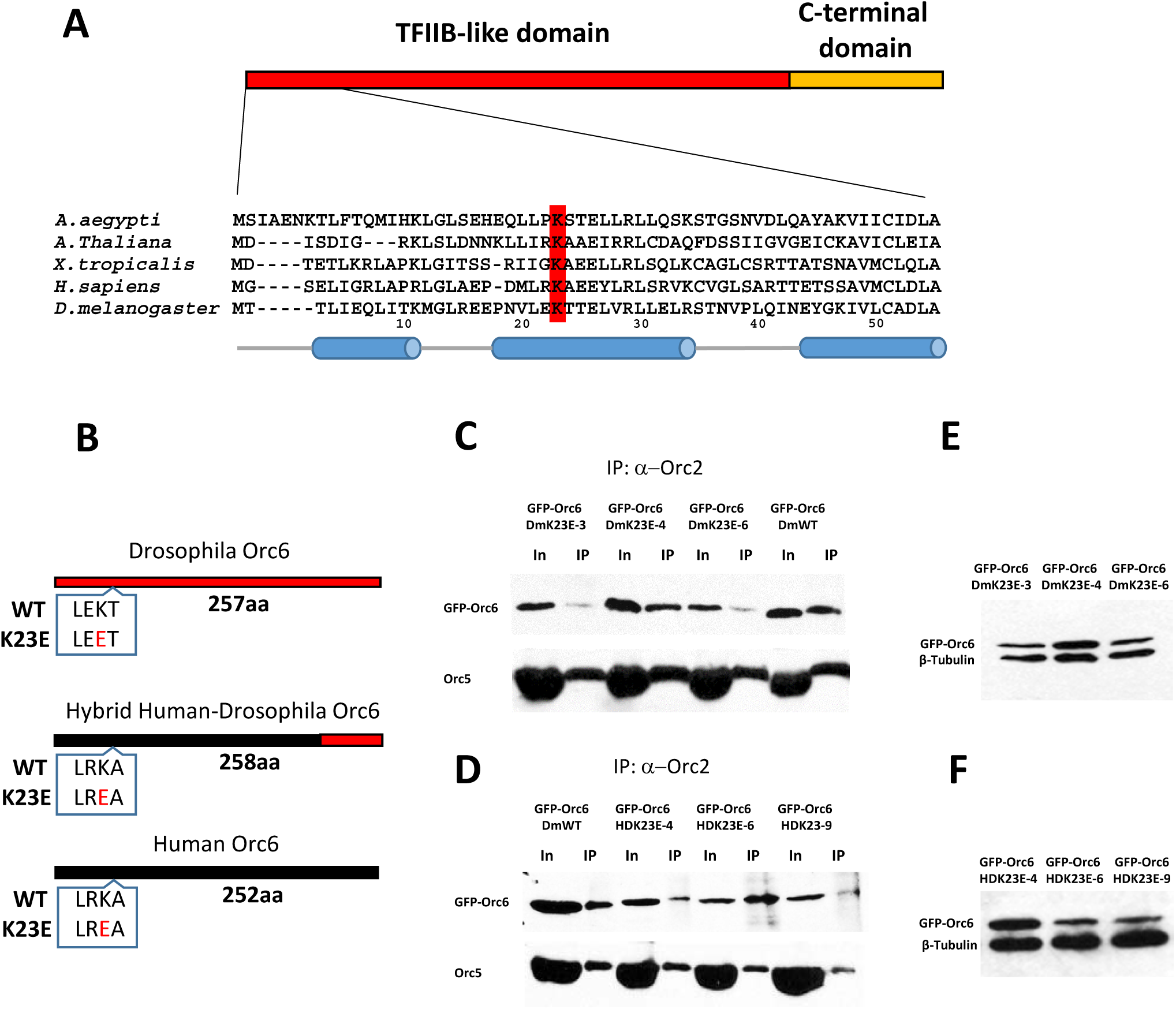
Orc6 mutants used in rescue experiments. **(A)** Amino acid sequence alignment of Orc6 proteins from different species. Conserved lysine at position 23 (K23) is indicated in red. Predicted helix structures in *Drosophila* and human proteins are shown in cylinders. **(B)** Positions of the K23E mutation are shown for Orc6 proteins used in this study. Mutation is colored in red. The hybrid human-*Drosophila* Orc6-HD protein contains first 206 amino acids of human Orc6 which includes TFIIB-like domain. **(C)** Immunoprecipitation (IP) of ORC complex using Orc2 antibodies from *Drosophila* Orc6 K23E mutant fly ovaries. IPs from three independent transgenic fly stocks are shown: K23E-3, K23E-4, K23E-6. **(D)** IP of ORC complex using Orc2 antibodies from hybrid human-*Drosophila* K23E mutant fly ovaries. Three independent transgenic fly stocks are shown: HDK23E-4, HDK23E-6, HDK23E-9. **(E**, **F)** The expression levels of mutant transgenic Orc6 proteins relative to tubulin.

The mutation in Orc6 leading to MGS (Y232S) was described before (Bicknell et al. 2011a), (de Munnik et al. 2012a), but it was located within C-terminal domain of the protein. In our earlier analysis of Orc6 functions *in vivo*, C-terminal Orc6 deletion (Orc6-200) as well as mutations of the conserved amino acids in the C-terminus of Orc6 D224A/Y225A and W228A/K229A also resulted in third instar lethality associated with defects in DNA replication and chromosome abnormalities (Balasov et al. 2009). Similar phenotypes were observed in our *Drosophila* model of the Meier-Gorlin Syndrome based on the conserved tyrosine to serine mutation (Y225S) in the C-terminal domain of Orc6 (Balasov et al. 2015). The molecular analyses of the mutation revealed that it disrupted the interaction between Orc6 and the rest of the ORC through the Orc3 subunit resulting in defects in pre-RC formation and MCM recruitment (Bleichert et al. 2013), (Balasov et al. 2015).

We asked a question what was a molecular basis of the MGS phenotype resulting from K23E mutation in the N-terminal domain of Orc6. Since only *Drosophila* Orc6 wild type and hybrid Orc6-HD resulted in viable adults we incorporated K23E mutation into *Drosophila* Orc6 gene and human N-terminal domain of Orc6-HD hybrid construct **(Figure 4B)**. We wanted to analyze the effect of this mutation for both *Drosophila* and human Orc6 proteins using “humanized” model of MGS in *Drosophila*. Orc6– DmK23E and Orc6-HDK23E constructs were introduced into flies carrying homozygous deletion of *orc6* gene. Three independent transgenic fly stocks were set up for each mutant.

First, we immunoprecipitated ORC complex using Orc2 antibodies from obtained transgenic flies and found that Orc6-DmK23E and Orc6-HDK23E proteins were both present in the corresponding complexes (**Figure 4C and 4D**). We noticed that Orc6-DmK23E co-immunoprecipitated better with the rest of the ORC complex when its expression level was higher (**Figure 4C and 4E**). Remarkably, stock Orc6-DmK23E-4 had the highest expression level of Orc6-DmK23E protein compared to others and promoted flies to the adult stage (**Figure 4E and Figure 5**). The amount of immunoprecipitated Orc6-HDK23E protein did not show noticeable correlation with the expression level of mutant proteins (**Figure 4D and 4F**) but the protein was always present in the IP material indicating that interaction of Orc6 carrying K23E substitution with core ORC was not affected by the mutation. It was shown before that metazoan Orc6 associated with ORC via its Orc3 subunit (Bleichert et al. 2013). Therefore, we tested whether K23E mutation would disrupt a direct interaction between Orc6 and Orc3 subunits. As shown in **Supplementary Figure 2C** mutant Orc6 protein carrying K23E mutation was able to pull down Orc3 in the *in vitro* transcription/translation reactions as well as the wild type *Drosophila* Orc6. We conclude that K23E mutation in Orc6 does not affect its association with the core ORC.

**Figure 5.**
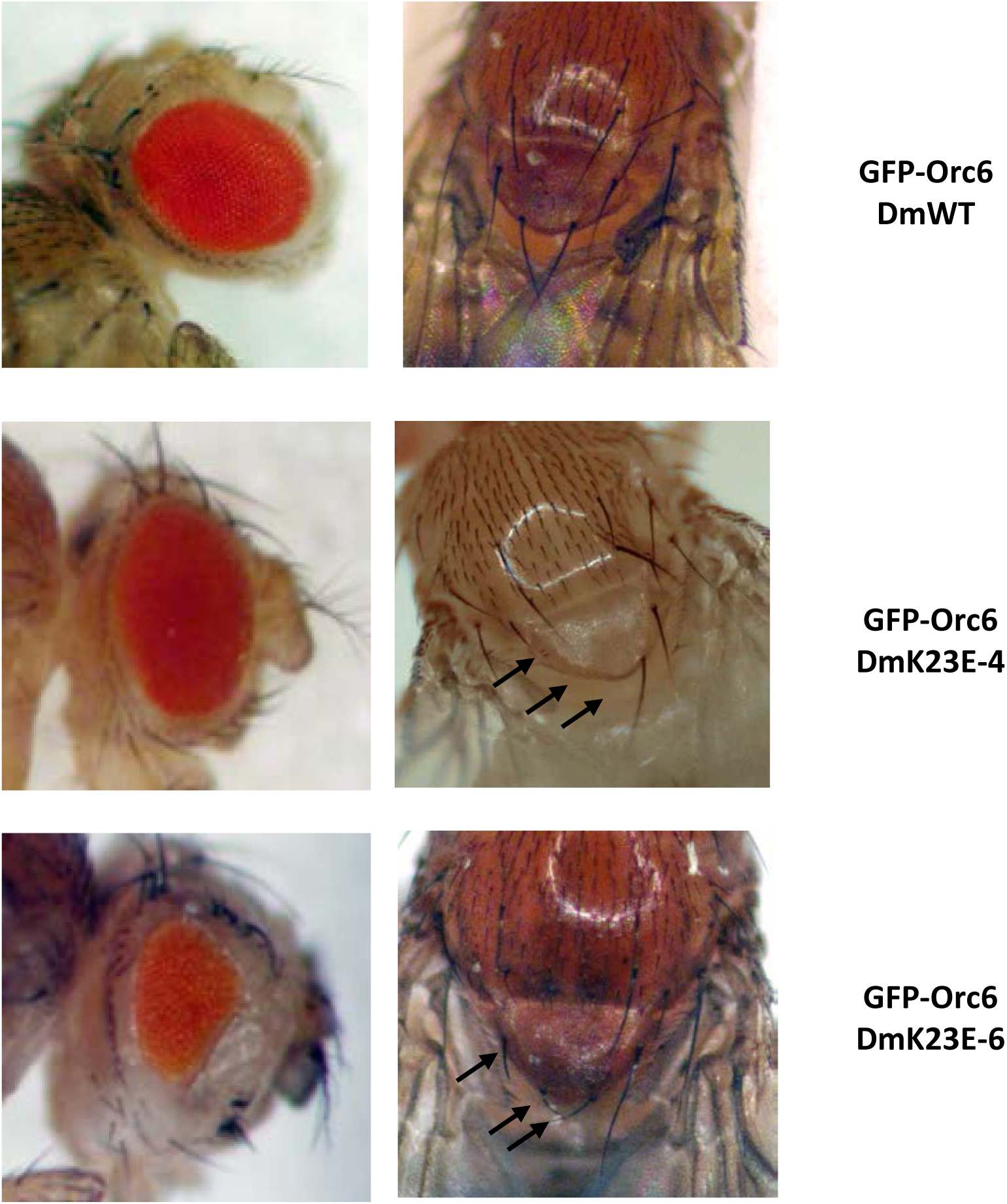
Phenotypes of *orc6* deletion flies rescued with Orc6-DmK23E. Flies carrying GFP-Orc6DmK23E-6 show reduced eye facets number and multiple defects of scutellar bristles. GFP-Orc6DmK23E-4 mutant flies have less severe phenotype: eye size and facets number are normal, however many flies have missing or defective scutellar bristles.

Second, we analyzed mutant phenotypes and found that only stock Orc6-DmK23E-4 resulted in viable adult flies majority of which were similar to the wild type. Minor defects (missing scutellar bristles) appeared in some flies (**Figure 5**). Another stock Orc6-DmK23E-6 produced very few homozygous adults. All of them had defective scutellar bristles and reduced eyes size (**Figure 5**). Eye of *Drosophila* is the well-established model to study proliferation defects. Compound eye consist of uniform regular facets and final number of them is determined during several mitotic divisions in third instar larvae. Any delay or disturbances in course of divisions result in reduced size and abnormal shape of adult eye. This phenotype was consistent with a disrupted Orc6 function in replication and cell cycle progression. Another interesting observed phenotype was inability of mutant flies to fly. Rescued adult flies carrying Orc6-DmK23E could not fly but were able to jump and walk. Previously we reported that flies with C-terminal Orc6 MGS mutation (Y225S) also missed scutellar bristles and were flightless (Balasov et al. 2015). The observed bristles defect and flightless phenotype could be a common “clinical” appearance of Meier-Gorlin syndrome in *Drosophila*.

To test if K23E mutation affects DNA replication we analyzed BrdU incorporation in fly stocks expressing mutant proteins Orc6-DmK23E and Orc6-HDK23E. Both mutants showed reduced BrdU incorporation in larval brain compared to wild type *Drosophila* or hybrid transposons (**Figure 2E and 2F**). Mitotic chromosomes in all six transgenic stocks looked fragmented and polyploid (**Figure 3B and 3C**). The strain expressing Orc6-DmK23E-4 displayed two different karyotypes; normal (**Figure 3B**, **middle left**) when all four chromosomes could be identified and nearly normal (**Figure 3B**, **middle right**) when all chromosomes could be identified but also additional pieces or broken ends were present. Since Orc6-DmK23E-4 flies were able to proceed to adult stage, we concluded that appeared chromosome defects were minor and not a life threatening in *Drosophila*. The minor chromosome defects were also reported for the human patient carrying K23E mutation (Li et al. 2017).

In our previous study describing fly model of MGS based on Orc6 Y225S mutation (Balasov et al. 2015)), we showed that the elevated expression of Orc6 Y225S transgene rescued *orc6* deletion phenotype. We asked, if it would be the case for K23E mutation. Similar to Y225S Orc6 mutant all K23E *orc6* transgenes have UAS promoter upstream of native *orc6* promoter. UAS promoter allows boosting expression by crossing transgenic flies with flies bearing *tubulin* promoter – driven GAL4 (see Material and Methods). As summarized in **Table 1**, elevated expression of both *Drosophila* and human-*Drosophila* hybrid transgenes carrying K23E mutation rescued lethality associated with Orc6 deletion. Remarkably, many of the rescued flies also had defects of the scutellar bristles and were not able to fly similar to the phenotypes observed in the fly model of MGS based on Orc6 Y225S mutation (Balasov et al. 2015). The expression of Orc6-HsK23E under either promoter did not rescue viability or any phenotypes associated with the *orc6* deficiency (**Table 1**).

**Table 1.**
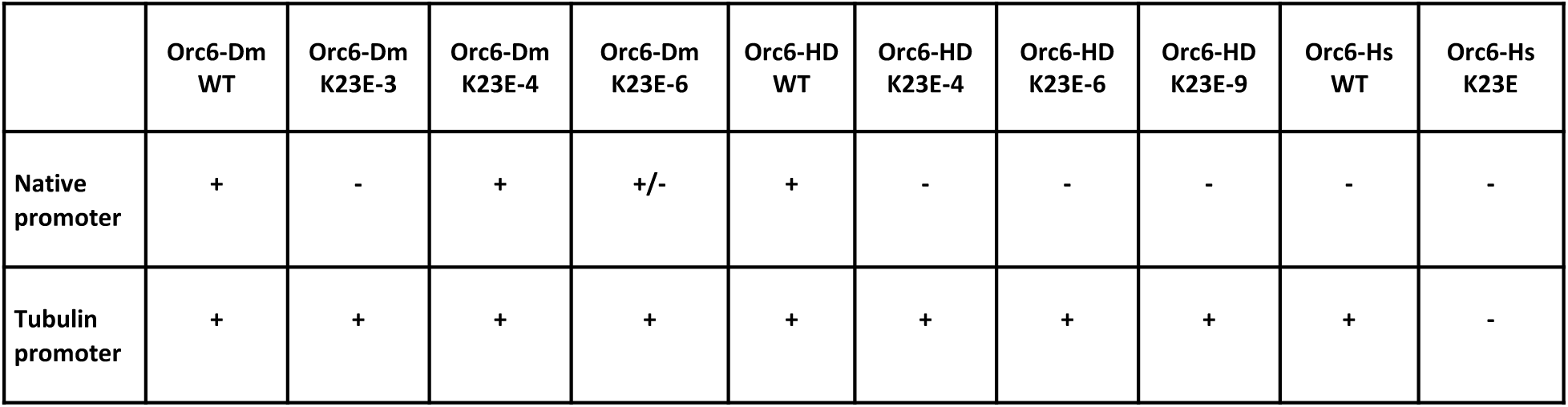
Rescue of the *orc6* deletion mutant with Orc6 K23E transgenes under control of native or tubulin promoter.

Overall, we concluded that both MGS mutations resulted in reduced DNA replication and chromosome defects, manifesting in similar phenotypes; however, the molecular mechanisms might be different. Mutation Y225S impedes Orc6 binding to ORC complex (Bleichert et al. 2013), (Balasov et al. 2015). Mutation K23E resides in the N-terminal domain of Orc6 important for DNA binding, therefore, next, we investigated the ability of *Drosophila*, human and hybrid Orc6 proteins carrying K23E mutation to bind DNA.

### 3. K23E mutations disrupt DNA binding ability of Orc6 protein

Six recombinant proteins were purified using *E.coli* expression system, *Drosophila* (Orc6-Dm-WT and Orc6-DmK23E), hybrid human/*Drosophila* (Orc6-HD and Orc6-HDK23E) and human (Orc6-Hs-WT and Orc6-HsK23E) (**Supplementary Figure 2A**). As shown in **Figure 6A**, both *Drosophila* Orc6 wild type protein and Orc6 carrying K23E mutation were able to bind *in vitro* with the ori-β DNA fragment derived from the origin of DNA replication associated with a chorion gene cluster. Binding was very tight with the majority of the labeled probe bound by the protein, as we observed before (Balasov et al. 2007) suggesting the presence of multiple binding sites for Orc6 within the bound ori-β fragments. However, the mobility of DNA-protein complex changed significantly when Orc6-K23E was used in the reaction. The mutant complex migrated faster compared to the wild type protein probably due to a reduced template occupancy resulting from K23E mutation. Increasing the concentration of the mutant protein in EMSA reactions only slightly improved the DNA binding of Orc6-K23E protein (**Supplementary figure 2B**).

**Figure 6.**
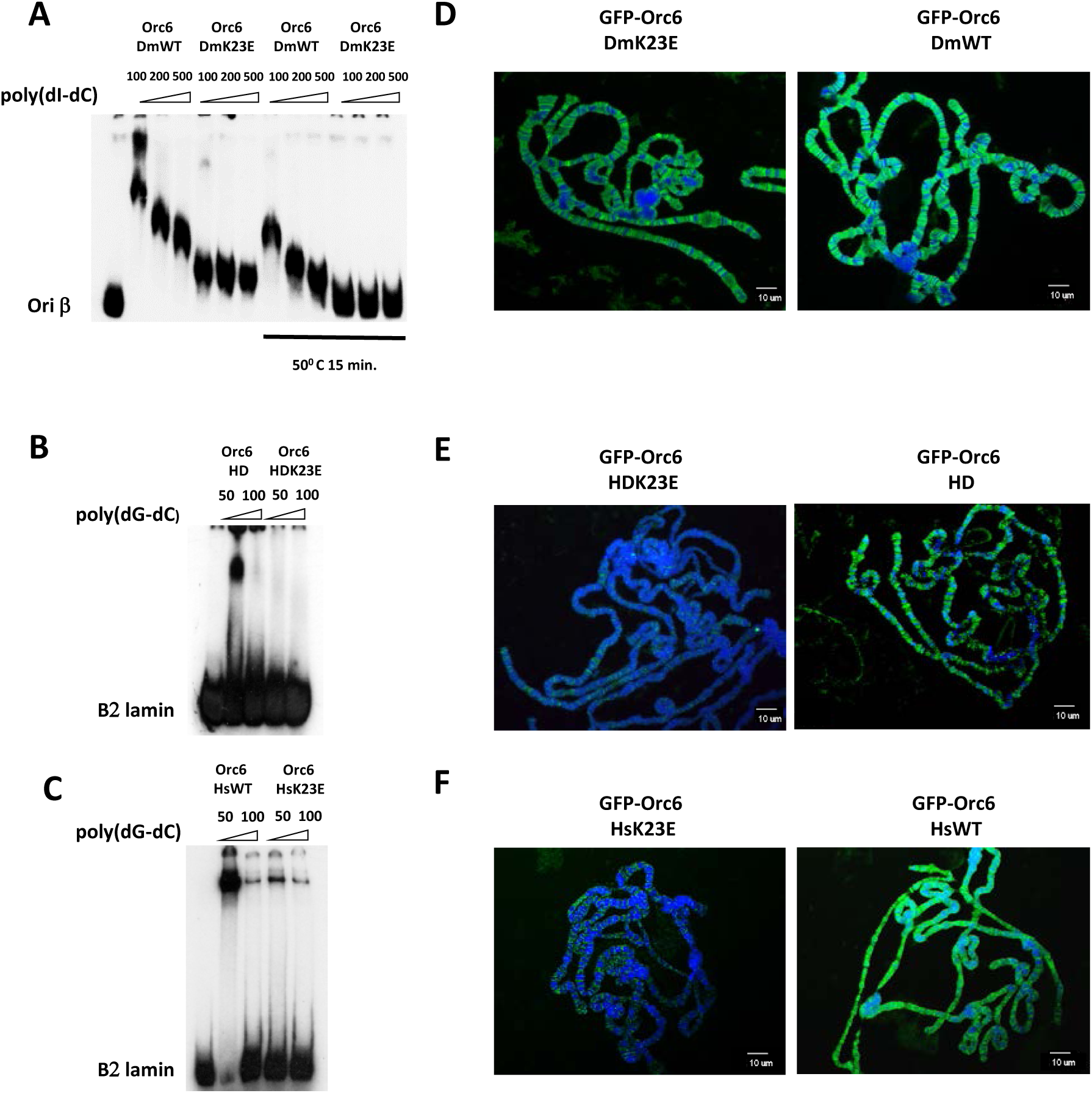
DNA and chromosome binding ability of hybrid and mutant Orc6. Electrophoretic mobility shift assay (EMSA) and chromosome binding of **(A) (D)** *Drosophila* Orc6 wild type and K23E mutant; **(B**, **E)** hybrid Orc6 HD wild type and Orc6 HD-K23E mutant proteins; **(C**, **F)** Human Orc6 wild type and K23E mutant. ‘50°C 15 min.’ – indicates that protein probe was heat-treated before adding to the reaction. Amount of competitor poly(dI-dC) and poly(dG-dC) is shown in nanograms. Polytene chromosomes isolated from salivary glands expressing GFP-tagged Orc6 were immunostained with anti-GFP antibodies, DNA was stained with DAPI.

When K23E mutation was introduced into human Orc6 protein or into a hybrid protein Orc6-HD containing human N-terminal TFIIB like domain, the overall DNA binding ability of the resulting mutant proteins was also significantly diminished (**Figure 6C and 6B**). We would like to point out that the DNA binding pattern for human Orc6 is different from *Drosophila* protein with the majority of the DNA-protein complex migrating at the top of the EMSA gel. This difference between fly and human Orc6 binding to DNA was reported before (Balasov et al. 2007), (Liu et al. 2011).

To visualize the Orc6 mutants binding *in vivo*, we used GFP-Orc6 bearing fly stocks described in Materials and Methods. The UAS promoter in the construct allows for GAL-4 induced expression using the GAL-4/UAS binary system (Brand and Perrimon 1993), (Duffy 2002). GFP-Orc6 expression was induced in salivary glands of third instar larvae to test chromosome binding of Orc6 mutants. Nuclei of *Drosophila* salivary glands contain polytene chromosomes that can be easily visualized with microscopy. GFP-fused Orc6-Dm-WT and Orc6-DmK23E (**Figure 6D**) were found to be associated with polytene chromosomes with Orc6-DmK23E showing slightly reduced binding as compared to wild type. In contrast, Orc6-HDK23E and Orc6-HsK23E mutants showed significantly reduced DNA binding and association with chromosomes (**Figure 6E and 6F**) correlated with the EMSA DNA binding experiments shown in **Figure 6B and 6C.** These data are in agreement with the survival pattern of corresponding fly strains (**Table 1**).

### 4. K23E mutation results in the instability of the *Drosophila* but not human Orc6

Often mutations lead to a protein unfolding that may affect its functions, such as DNA binding. Therefore, we tested the effect of K23E mutation on protein stability. We measured tryptophan fluorescence of *Drosophila* and human Orc6 proteins at the increasing temperatures and compared them with mutants carrying K23E mutation. As presented in **Figure 7A**, K23E mutation in human Orc6 did not change a stability of the mutant protein relative to wild type at all tested temperatures, however, *Drosophila* Orc6 carrying K23E was more susceptible to the heat and began unfolding at the temperatures over 35°C suggesting that the mutation causes an instability in fly protein. Interestingly, fluorescent intensity of *Drosophila* mutant relative to wild type was significantly lower even at 25°C (**Figure 7B**). This indicates that some part of the mutant protein is already unfolded at temperatures optimal for *Drosophila* development. Interestingly, *Drosophila* Orc6 and hybrid Orc6 proteins with *Drosophila* C-terminus were more prone to degradation during purification compared to the human Orc6 proteins (**Supplementary Figure 2A and data not shown**). We hypothesized that thermal instability may further impact DNA binding ability of *Drosophila* Orc6. To test this hypothesis we repeated electro mobility shift assay with wild type and K23E mutant protein probe being heat shocked at 50°C for 15 minutes. Mutant Orc6 did not show any DNA binding under these conditions whereas the wild type Orc6/DNA complex migrated faster compared to the protein not subjected to heat treatment (**Figure 6A and Supplementary Figure 2B**). It is possible that the instability of Orc6-DmK23E protein together with a loss of DNA binding ability may contribute to the MGS phenotype in *Drosophila*.

**Figure 7.**
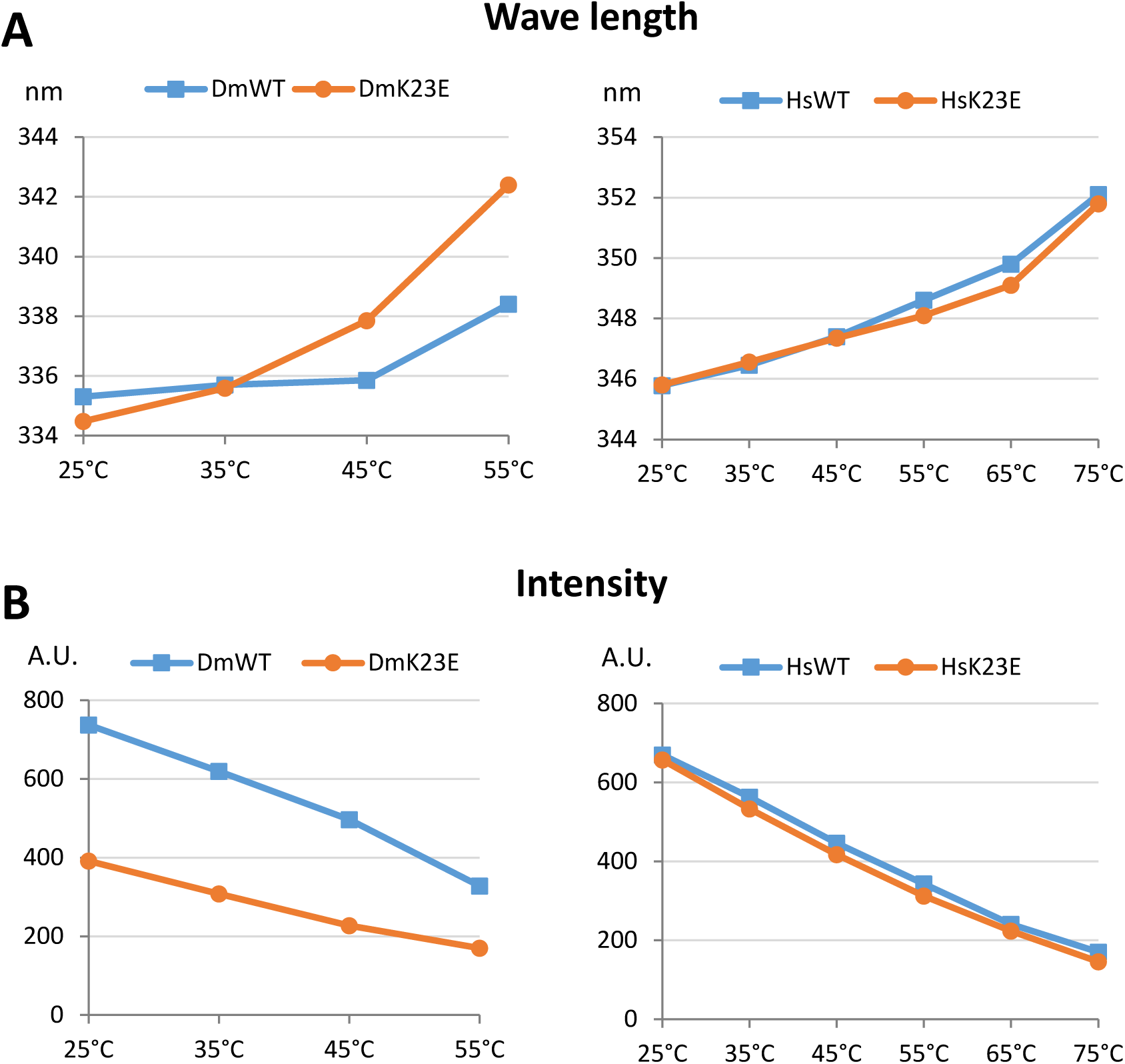
Measuring of *Drosophila* and human Orc6 protein stability by fluorescence spectroscopy. **(A)** Emission maxima shifts at gradually increasing temperatures. **(B)** Absolute intensity at each temperature point. DmWT and DmK23E – *Drosophila* Orc6 wild type and Orc6-K23E mutant; HsWT and HsK23E-human Orc6 wild type and Orc6 Hs-K23E mutant.

### 5. Elevated expression of the wild type human Orc6 in *Drosophila* rescues a lethality associated with a deletion of the *orc6* gene

In our previous work (Balasov et al. 2015) we found that a *Drosophila* C-terminal MGS mutation Y225S was lethal in flies despite the mild MGS clinical appearance in humans. Importantly, human Orc6 is loosely associated with the core ORC (Vashee et al. 2001), (Vashee et al. 2003), (Ranjan and Gossen 2006), whereas *Drosophila* Orc6 associates with core ORC much more tightly (Gossen et al. 1995), (Chesnokov et al. 1999), (Chesnokov et al. 2001). This might explain why the Y225S MGS mutation in Orc6 has more dramatic effect for survival of *Drosophila* compared to corresponding Y232S mutation in humans. We also found that elevated expression of mutant protein carrying Y225S mutation rescued lethality, restored normal karyotype and allowed detection of Orc6 with the rest of ORC complex (Balasov et al. 2015). The expression of the wild type human Orc6 under native *orc6* promoter could not rescue a lethality associated with *Drosophila orc6* gene deletion (**Table 1**). We asked if overexpression of the human Orc6 would rescue *orc6* deficiency in *Drosophila*. Again, using GAL4/UAS binary system, we boosted the expression of human wild type and K23E mutant in flies. We found that the elevated expression of the wild type human Orc6 was able to rescue flies to viability (**Table 1**) and restored normal karyotype (**Figure 8A**). We also observed human Orc6 with other ORC subunits after immunoprecipitation with anti-Orc2 antibodies (**Figure 8B**). Remarkably, rescued adult flies displayed upheld wing phenotype (**Figure 8 C**, **D**) and were not able to fly (**tub-Orc6-HsWT movie file**) similar to the *Drosophila* C-terminal MGS mutation Y225S flies (Balasov et al. 2015). However, when K23E mutation was introduced into the human Orc6 the resulting mutant Orc6-Hs-K23E gene was not able to rescue *orc6* deficient flies to viability even under overexpressed condition and no improvements were observed during development of the flies (**Figure 8A**, **lower row**). Wild type human Orc6 loosely associates with ORC complex (Vashee et al. 2001), (Vashee et al. 2003), (Ranjan and Gossen 2006) somewhat mimicking the effect of Y225S mutation in *Drosophila* Orc6. As a result, Orc6-Hs-K23E protein appears to be defective in both DNA binding (**Figure 6**, **C and F**) and in the interaction with a core ORC, therefore it was not possible to rescue a lethality regardless of the protein expression level (**Table 1**).

**Figure 8.**
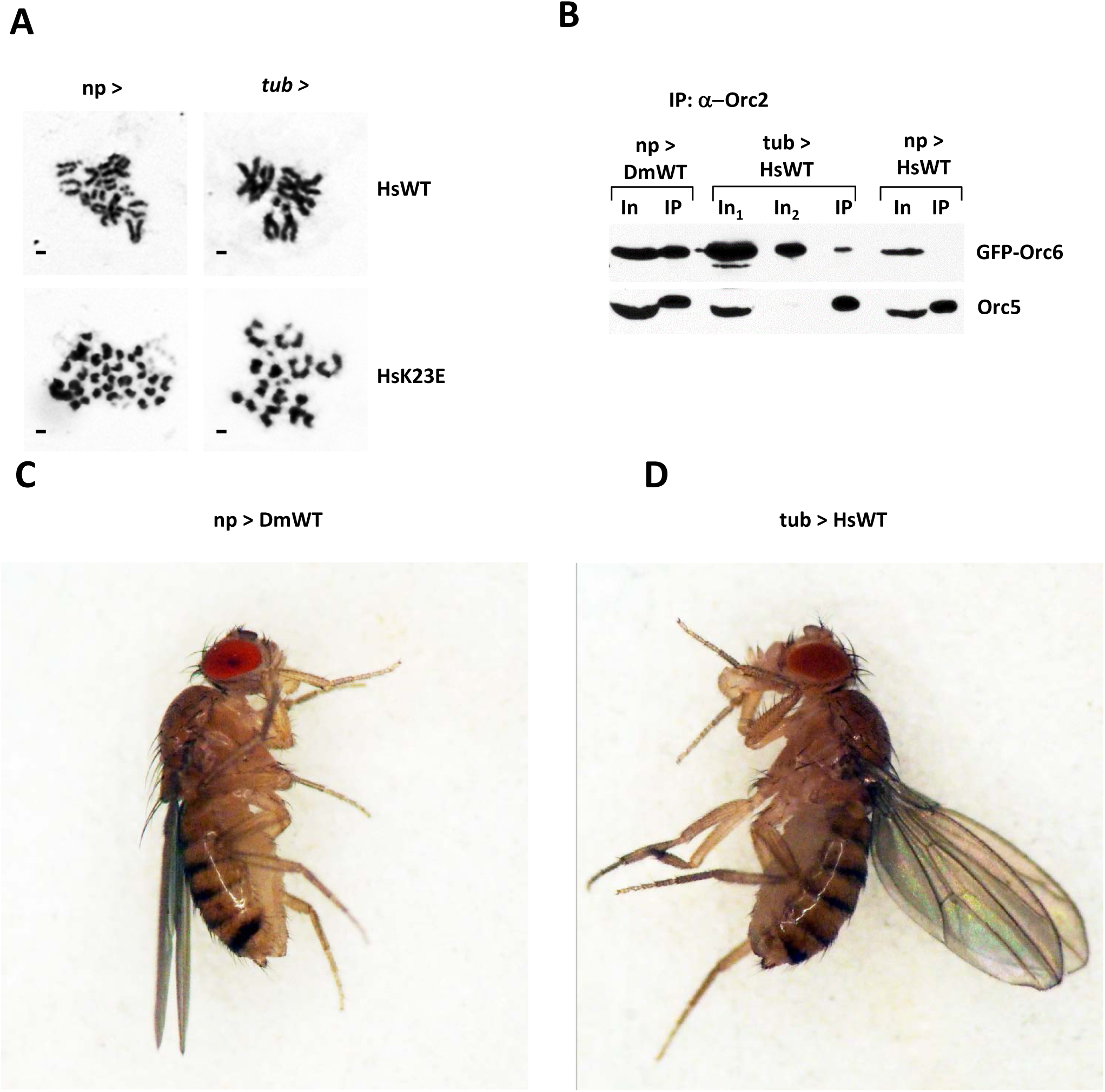
Rescue of *orc6* deletion flies with human Orc6. **(A)** Mitoses in neuroblasts of *orc6* deletion mutant rescued with human Orc6 wild type or with K23E MGS mutation. Expression of transgenes was driven with either native *orc6* promoter (np >) or with strong constitutive *tubulin* promoter (tub >). **(B)** Immunoprecipitation of the ORC complex with anti-Orc2 antibodies from ovaries expressing GFP tagged Orc6 transgenes. In1 – 1/10, In2 – 1/50 of total IP reaction. **(C)** Adult flies rescued with wild type *Drosophila* Orc6, **(D)** Adult flies rescued with *tubulin* promoter driven overexpression of wild type human Orc6.

### 6. Microarray analysis of the MGS mutation in *Drosophila*

In our previous study we found that flies carrying Y225S mutation had rough spots in the eye, irregular hair patterns, missing bristles – phenotypes, possibly associated with minor cell polarity defects (Balasov et al. 2015). These phenotypes could indicate potential role of Orc6 in other processes or pathways apart from replication. To investigate this possibility further, we conducted a microarray analysis of gene expression to identify the genes/pathways which are affected by the MGS mutations. Orc6 carrying Y225S MGS mutation rescued adult flies only when overexpressed, therefore *orc6*^*35*^*/orc6*^*35*^;*Orc6-Y225S/tubP-GAL4* adults were subjected to the microarray analysis against *orc6*^*35*^*/orc6*^*35*^;*Orc6-WT/tubP-GAL4* adults. From the total analyzed 13557 genes, 1446 genes were downregulated more than 2 times and 1363 genes were upregulated more than 2 times. Gene ontology analysis revealed downregulated DNA replication, DNA repair and cell proliferation processes in top 10 scores (**Supplementary table 1**). One of Orc6-K23E MGS mutant transgenes rescued flies to the viability without overexpression, therefore, *orc6*^*35*^*/orc6*^*35*^;*Orc6-K23E-4/Orc6-K23E-4* were analyzed and compared to *orc6*^*35*^*/orc6*^*35*^;*Orc6-WT/Orc6-WT*. In this case, only 33 genes displayed more than 2 times downregulation and 25 genes – more than 2 times upregulation. This result is consistent with a relatively mild phenotype associated with K23E mutation compared to Y225S mutation. Next, we filtered up and down regulated genes in both mutants for common genes. The end results consisted of two short lists of 9 downregulated genes and 14 upregulated genes (**Supplementary table 1**). We have analyzed the expression patterns of genes based on modeENCODE RNA-seq data (flybase.org) and found that all 9 downregulated genes had maximum expression level in ovaries. Ovary is the organ where the majority of the replication events happens in adult flies, therefore they are more sensitive to replication defects. The 14 upregulated genes showed high expression in head and digestive system. Most of them encoded products with endopeptidase and ubiquitin-transferase activities. We hypothesize that the elevated expression of these genes could reflect degradation/utilization processes in tissues where proliferation defects resulted in cell damage. Overall microarray analysis did not reveal any additional genes/pathways affected by MGS mutations with the exception of replication and proliferation.

### 7. The analysis of flightless phenotype

The flightless phenotype is a characteristic of all analyzed Orc6-based MGS mutants in *Drosophila* including previously described Y225S (Balasov et al. 2015) and a new K23E mutations. In our current study GAL4>UAS K23E boost rescued the lethality and bristle defects but did not restore the ability to fly. We assayed flight performance of the GAL4>UAS K23E boosted mutants by dropping flies individually into 90cm cylinder. Flies expressing *Drosophila* Orc6-WT immediately escaped the cylinder or landed on the wall and flew away in seconds. In contrast, all mutants landed on the bottom of the cylinder, they did not try to escape and were easy to collect. Interestingly, the flies rescued with human Orc6 were not able to fly either (**tub-Orc6-HsWT movie file**).

To understand the cellular mechanism of flightless phenotype we prepared histological analysis of the major thoracic muscles. In *Drosophila*, large indirect flight muscles (IFMs) mediate flight by contraction and expansion of the thoracic cuticle. IFMs are constituted of six dorsal longitudinal muscles (DLMs) and seven dorsoventral muscles (DVMs) (Bernard et al. 2003). Therefore we analyzed histological transverse section of IFMs for signs of degeneration or atrophy. Light microscopy revealed that all large bundled muscle fibers group were present in MGS mutant (**Supplementary figure 3)**. An inability to fly might also result from defects in IFMs innervation. To test this we induced RNAi of *orc6* specifically in motor neurons. Remarkably, the obtained flies were not able to fly. This experiment suggests that Orc6 might participate in motor neurons development during metamorphosis.

## Discussion

Meier-Gorlin syndrome (MGS) is a rare human disease associated with microcephaly, short stature and multiple developmental defects (de Munnik et al. 2012b), (de Munnik et al. 2012a), (Bicknell et al. 2011a), (Kerzendorfer et al. 2013). Mutations in a number of factors involved in DNA replication have been found to be causative for this disease (Bicknell et al. 2011a), (Bicknell et al. 2011b), (Guernsey et al. 2011), (Fenwick et al. 2016), (Vetro et al. 2017), (Burrage et al. 2015), (McDaniel et al. 2020). Several mutations causing MGS were found in ORC subunits including Orc6 (Bicknell et al. 2011a), (Bicknell et al. 2011b), (Guernsey et al. 2011). In humans, Y232 to S substitution in Orc6 corresponds to a mild clinical appearance of MGS (Bicknell et al. 2011a), (de Munnik et al. 2012a), however, corresponding mutation in *Drosophila* (Y225S) is lethal and molecular and cell analysis revealed significantly reduced DNA replication and chromosome fragmentation in cells and tissues (Balasov et al. 2015). Importantly, the flies carrying Orc6-Y225S mutation can be rescued by the elevated expression of the transgene (Balasov et al. 2015).

The second case of MGS related to human Orc6 is a homozygous deleterious mutation within the gene. Small deletion c.602-605delAGAA generated stop codon after 202 amino acid (Shalev et al. 2015). The severity of an abnormal embryological development coincided with a lethal embryonic phenotype in *Drosophila* for Orc6 having only first 200 amino acids (Balasov et al. 2009). The elevated expression of this mutant protein did not rescue a lethal phenotype of flies (Balasov et al. 2015).

The third case of MGS was recently reported and connected with a mutation in the conserved lysine at position 23 resulting in Lys23 to Glu mutation (Li et al. 2017). This mutation caused a range of phenotypes in *Drosophila* from lethality to relatively normal adults and the severity of the observed phenotypes correlated with the expression level of the mutant proteins. This newly described K23E mutation localizes in the N-terminus of the protein. Not surprisingly, biochemical and cytological analysis revealed that K23E mutation in both *Drosophila* and human Orc6 reduced DNA binding in gel shift experiments. *In vivo* this mutation leads to inability of Orc6-HsK23E or Orc6-HDK23E to bind with chromosomes but the chromosome association of Orc6-DmK23E was not significantly affected. This observation coincided with the survival rates as flies carrying Orc6-DmK23E often were able to progress to the adulthood. Interestingly, K23E mutation leads to the instability of the fly protein at the elevated temperatures with a difference noticeable already at 37°C. However, in the case of human protein, the same mutation did not change protein conformation compared to the wild type even at the higher temperatures. Normal body temperature for humans at 37°C is lethal for *Drosophila*, suggesting that evolutionary changes occurred in warm blooded animals to withstand thermal challenges otherwise lethal for insects. In summary, the two known Meier-Gorlin Syndrome mutations in Orc6 reside in different functional domains of the protein and result in either impaired DNA binding (K23E) by Orc6 or a loss of the protein association with the core ORC (Y232S). The consequences of both mutations include the reduced amounts of hexameric ORC on DNA, an impaired pre-RC formation and fewer origin firing. This in turn leads to the replication, proliferation and development defects manifesting in similar phenotypical traits in both cases.

*Drosophila* and human Orc6 proteins have only 28% of sequence identity but are structurally similar and critical for the initiation of DNA replication (Duncker et al. 2009). The metazoan Orc6 consists of two domains, a larger N-terminal domain (∼200 amino acids) which carries homology with TFIIB transcription factor and is important for DNA binding (Chesnokov et al. 2003), (Balasov et al. 2007), (Liu et al. 2011) and a shorter C-terminal domain important for the function of Orc6 in cytokinesis (Prasanth et al. 2002), (Chesnokov et al. 2003), (Huijbregts et al. 2009), (Bernal and Venkitaraman 2011), (Akhmetova et al. 2015), as well as for the interaction of Orc6 with core ORC (Bleichert et al. 2013). Full length human Orc6 could not rescue a lethality associated with a loss of the fly gene when expressed under native *orc6* promoter. Therefore, we created the hybrid Orc6 protein containing human N-terminal domain and *Drosophila* C-terminus. This transgene rescued *orc6* deletion flies to viability and they were undistinguishable from the wild type animals. This indicates that N-terminal TFIIB-like domain of both proteins involved in the same conserved functions between organisms. On the other hand, C-terminus of both *Drosophila* and human Orc6s contains conservative motifs responsible for association with ORC complex. Both flies and humans with C-terminal truncations do not survive to adult stage (Balasov et al. 2009), (Shalev et al. 2015). However, point mutation (Y232S) in human Orc6 manifests in a mild postnatal phenotype (Bicknell et al. 2011a), (de Munnik et al. 2012a), while the corresponding *Drosophila* Y225S MGS mutant shows no difference from lethal *orc6* deletion. It is known that human Orc6 is loosely associated with the core ORC (Vashee et al. 2001), (Vashee et al. 2003), (Ranjan and Gossen 2006), whereas *Drosophila* Orc6 associates with other ORC subunits significantly more tightly (Gossen et al. 1995), (Chesnokov et al. 1999), (Chesnokov et al. 2001). This might explain why the Y225S MGS mutation has more dramatic effect for survival of *Drosophila* than corresponding MGS (Y232S) mutation in humans.

In our earlier study (Balasov et al. 2015) we found that an elevated expression of the fly protein carrying Y225S mutation rescued third instar lethality, restores normal karyotype and mutant protein was detected on DNA with the rest of ORC complex. Similarly, here we found that the elevated expression of human Orc6 allowed the formation of the functional six-subunit ORC on DNA and rescued flies carrying *orc6* deletion. In some way human Orc6 in fly system mimics the effect of the C-terminal MGS mutation in *Drosophila* Orc6. In both cases the interaction of the proteins with core ORC is impaired resulting in similar phenotypes between flies carrying either *Drosophila* Orc6 with MGS or the human protein as a sole source of Orc6.

In this study we showed that hybrid protein Orc6-HD containing intact human N-terminal TFIIB like domain (∼80% of the protein length) and *Drosophila* C-terminus tightly associated with ORC and rescued Orc6 deficient flies to viable adults phenotypically undistinguishable from wild type animals. This hybrid approach revealed the importance of evolutionary conserved and variable domains of Orc6 protein and allowed the studies of human protein functions and the analysis of the critical amino acids in live animal heterologous system. We believe that hybrid approach not only open a broad avenue to study new Orc6 mutations for medical and general science purposes but might be useful in other humanized models. In summary, the humanized fly model presented in our studies has the unique advantage of being able to differentially test both fly, human and chimeric Orc6 proteins to reveal conserved and divergent features of the protein and its functions in the cells of metazoan organisms. The fly strains generated during this work will be useful in analyzing the functions of human protein in a convenient heterologous system. Specifically, our studies revealed novel insights into molecular mechanisms underlying MGS pathology and provide important clues about disease origin and development.

## MATERIALS AND METHODS

### Mutagenesis

The conservative lysine at position 23 was replaced with glutamic acid to create Meier-Gorlin mutation (K23E) following site-directed mutagenesis protocol (Agilent). *Drosophila*-Human and Human-*Drosophila* hybrids were designed by using a PCR technique. Orc6 mutant and hybrid genes were cloned in frame with GFP into the modified pUAST vector containing *UAS* promoter and *orc6* native promoter. All constructs were injected into *Drosophila w*^*1118*^ embryos (Model System Injections, Raleigh, NC) and individual transgenic strains were set up.

### Purification of recombinant Orc6

Wild type, mutant or hybrid Orc6 cDNAs were cloned into pET15b expression vector and transformed in BL21 *E.Coli* strain. Ni-NTA purified Orc6 proteins were further purified with HiTrap SP HP (GE Life Sciences) and Superdex 75 (GE Life Sciences) columns.

### Electrophoretic mobility shift assay (EMSA)

1 ng of P^32^ end labeled Human B2 lamin or *Drosophila* ori-β fragments were incubated with 300 ng of purified protein in reaction buffer (25mM Tris, pH 8, 60mM KCl, 5mM MgCl2, 0.1% NP-40, 0.12 mg/ml BSA, 10% glycerol) at room temperature for 30 min. 10 µl of each reaction was loaded on a 4% native polyacrylamide gel. Electrophoresis was performed at room temperature using TAE pH 8.3 for *Drosophila* Orc6 and pH 9.5 for human Orc6 as a running buffer. The gel was dried on Whatman paper and exposed to X-ray film.

### Fly stocks and rescue experiments

Following fly stocks containing endogenous *orc6*^*35*^ deletion and *GFP-orc6* fused transposon were set up:

1. *orc6*^*35*^*/Cy, GFP-orc6-wt* : wild type *Drosophila* Orc6
2. *orc6*^*35*^*/Cy, GFP-orc6-K23E:* K23E MGS *Drosophila* Orc6
3. *orc6*^*35*^*/Cy, GFP-orc6-Hs-wt:* wild type human Orc6
4. *orc6*^*35*^*/Cy, GFP-orc6-HsK23E:* K23E MGS type human Orc6
5. *orc6*^*35*^*/Cy, GFP-orc6-HD:* interspecies hybrid
6. *orc6*^*35*^*/Cy, GFP-orc6-DH:* interspecies hybrid
7. *orc6*^*35*^*/Cy, GFP-orc6-HDK23E:* : interspecies hybrid with K23E MGS mutation

Bloomington stock #5138 (*y,w; P{w*^*+mC*^*=tubP-GAL4}LL7/TM3,Sb,Ser)* expressing GAL4 ubiquitously under the control of the *alphaTub84B* promoter was used to design *orc6*^*35*^*/Cy; tub-GAL4/TM3,Sb* fly stocks for rescue experiments.

In rescue experiments with native promoter, progeny from heterozygous *orc6*^*35*^*/Cy; GFP-Orc6* was analyzed for the presence of *orc6*^*35*^*/orc6*^*35*^; *GFP-Orc6* adult flies. In rescue experiments with *alphaTub84 (tub)* promoter, females of the genotype *orc6*^*35*^*/Cy; GFP-Orc6* were crossed to males *orc6*^*35*^*/Cy; tub-GAL4/TM3,Sb*, and resulting progeny analyzed for the presence of the *orc6*^*35*^*/orc6*^*35*^;*GFP-Orc6/tub-GAL4* adults.

For Orc6 depletion in motor neurons we used Orc6 RNAi fly stock #HMJ22188 (NIG-FLY stock center, Japan) and motor neuron specific D42-GAL4 driver #8816 (Bloomington stock center).

### BrdU labeling and immunostaining of third-instar larval brains

Larval brains were soaked in PBS with 1µM BrdU for 30 min at 25°C. BrdU incorporation was detected by monoclonal antibodies (Becton Dickinson) following manufacturer protocol as described previously (Sullivan 2000).

### Immunostaining of polytene chromosomes

The flies bearing *GFP-orc6* transposon were crossed to Bloomington fly stock 6870 (w^1118^; *P{Sgs3-GAL4.PD}TP1)* expressing GAL4 under *Sgs3* promoter. *Sgs3* promoter drives GAL4 in the salivary glands of third-instar larvae. Salivary glands expressing different variants of GFP-Orc6 were dissected in PBS with 0.1% NP-40, transferred in fixing solution (2% formaldehyde, 45% acetic acid) for 1 min, squashed in 45% acetic acid and frozen in liquid nitrogen. The slides were desiccated in 96% alcohol and stored in 70% alcohol at −20°C. For immunofluorescence studies, slides were briefly washed in PBS with 0.1% NP-40 and incubated with primary antibodies (mouse anti-GFP (B2), Santa Cruz, #sc9996) diluted in 10% goat antiserum for 2 hours in humid chamber. Secondary goat anti-mouse antibodies (Alexa fluor 488, Thermo/Fisher) visualized Orc6 on the polytene chromosomes.

### Immunoprecipitation (IP) from ovaries

20 freshly dissected ovaries were crushed in the glass homogenizer with 100µl of high-salt IP buffer (25mM Hepes pH 7.6, 12.5mM MgCl2, 100mM KCl, 0.1mM EDTA, 300mM NaCl, 0.01% Triton X-100) and extracted for 1 hour at 4°C with continuous rotation. Extract was centrifuged at 15000g for 15 minutes and supernatant was diluted 3 times with low-salt IP buffer (25mM Hepes 7.6, 12.5mM MgCl2, 100mM KCl, 0.1mM EDTA, 0.01% Triton X100). Protein A-Sepharose (BioVision #6501-5) and rabbit polyclonal antibodies against Orc2 subunit were added and incubated with supernatant overnight. Sepharose beads were washed 3 times with low-salt IP buffer and diluted in 10 µl of IP buffer. Samples were boiled in loading buffer, separated in 10% SDS-polyacrylamide gel and transferred on Immobilon-P membrane (Millipore #IPVH00010).

### Mitotic Chromosome Preparation

Preparation of mitoses was described previously (Lebedeva et al. 2000). Briefly, third instar larval neural ganglia were incubated in 0.075M KCl for 5 minutes, fixed in methanol with acetic acid (3:1) for 20 minutes and then dispersed in a drop of 50% propionic acid on a slide. Then slides were dried and stained with 5% Giemsa’s solution.

### Western blot of larval brains

Homozygous (*orc6*^*35*^*/orc6*^*35*^, *GFP-orc6-HD, orc6*^*35*^*/orc6*^*35*^, *GFP-orc6-DH* versus heterozygous (*orc6*^*35*^*/Cy-YFP, GFP-orc6-HD, orc6*^*35*^*/Cy-YFP, GFP-orc6-DH*) third instars larvae were selected based on fluorescence of the *Cy-YFP* balancer. Fresh dissected brains were incubated with 450mM NaCl, 0.5% NP-40 in PBS buffer for 1 h and samples were centrifuged 10 minutes at 10000g. Pellets were washed 3 times with 100mM NaCl, 0.5% NP-40 in PBS buffer and used for western blot. MCM complex was detected with rabbit polyclonal anti MCM4 and anti MCM5 antibodies.

### Fluorescence Spectroscopy

Purified Orc6 protein (wild type or mutant) was diluted in buffer (50mM NaH2PO4, pH 7.4, 100mM NaCl) to 10 ng/µl. 260 µl samples were loaded in quartz cuvette. Fluorescence measurements were performed in the VARIAN CARY Eclipse fluorescence spectrophotometer. The spectral widths of the excitation and the emission bands were 5 nm and 20 nm, respectively. Excitation was performed at 290 nm wavelength. Emission spectra were recorded from 300 to 400 nm in 1-nm steps. An emission spectrum of a buffer was subtracted from protein spectra.

### Microarray analysis

RNA was isolated from five days old females using ZR Tissue and Insect RNA MicroPrep™ (Zymo Research, #R2030). DNA was removed using TURBO™ DNase (Invitrogen, #AM2238) following manufacturer’s recommendations. cDNA was generated from 1 μg of total RNA using ProtoScript® II First Strand cDNA Synthesis Kit (New England Biolabs, #E6560).

Microarray experiment was conducted at the Boston University Microarray and Sequencing Resource Core Facility. Drosophila Gene 1.0 ST CEL files were normalized to produce gene-level expression values using the implementation of the Robust Multiarray Average (RMA) (Irizarry et al. 2003) in the *affy* package (version 1.48.0) (Gautier et al. 2004) included in the Bioconductor software suite (version 3.2) (Gentleman et al. 2004) and an Entrez Gene-specific probeset mapping (20.0.0) from the Molecular and Behavioral Neuroscience Institute (Brainarray) at the University of Michigan (Dai et al. 2005), (brainarray.mbni.med.umich.edu/Brainarray/Database/CustomCDF). Array quality was assessed by computing Relative Log Expression (RLE) and Normalized Unscaled Standard Error (NUSE) using the *affyPLM* package (version 1.46.0). Principal Component Analysis (PCA) was performed using the *prcomp* R function with expression values that had been normalized across all samples to a mean of zero and a standard deviation of one. Differential expression was assessed using the moderated (empirical Bayesian) *t* test implemented in the *limma* package (version 3.26.9) (i.e., creating simple linear models with *lmFit*, followed by empirical Bayesian adjustment with *eBayes*). Correction for multiple hypothesis testing was accomplished using the Benjamini-Hochberg false discovery rate (FDR). Human homologs of fly genes were identified using HomoloGene (version 68). All microarray analyses were performed using the R environment for statistical computing (version 3.2.0).

Gene Ontology terms analysis was conducted using DAVID Bioinformatics Resources 6.8 (david.ncifcrf.gov) (Huang da et al. 2009).

## Acknowledgements

We would like to thank Adam Gower from the Boston University Microarray and Sequencing Resource Core Facility. This work was supported by a grant from NIH to IC (GM121449).

## Author contributions

M.B., K.A. and I.C. designed and performed the experiments. M.B., K.A. and I.C. prepared the figures. I.C. wrote the manuscript with the help of M.B. and K.A. All authors approved the manuscript.

## Notes

### Competing Interest Statement

The authors have declared no competing interest.

