## Supplementary Figures and Table for "Humanized *Drosophila* model of the Meier-Gorlin syndrome reveals conserved and divergent features of the Orc6 protein"

### Supplementary figure 1

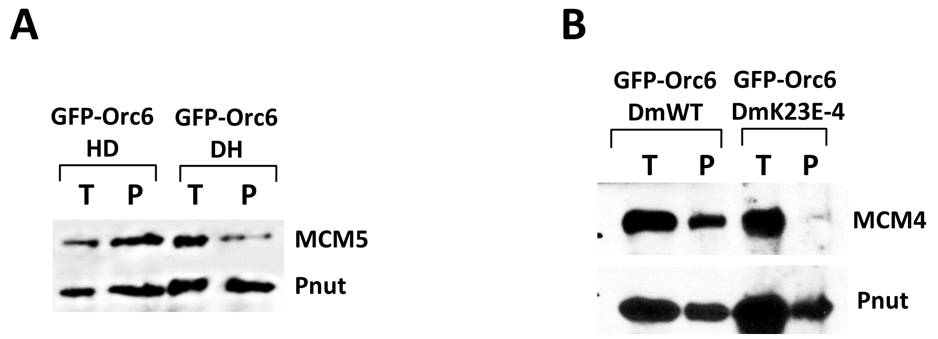


**P – insoluble pellet after salt extraction**

**T – total brain extract**

**MCM2–7 complex chromatin association. (A)** Chromatin association of Mcm5, a member of the MCM2–7 complex, is reduced in *orc6* deletion fly larvae expressing GFP-Orc6-DH but not GFP-Orc6-HD hybrid protein. Neural ganglia of homozygous, *orc6* deletion *Drosophila* larvae expressing GFP-Orc6 transgenes under native *orc6* promoter were isolated and subjected to salt extraction to solubilize cellular proteins. Insoluble and chromatin associated proteins were pelleted by centrifugation and analyzed by Western blotting using Mcm5 polyclonal antibodies. Pnut was used as a loading control. **(B)** Chromatin association of Mcm4, a member of the MCM2–7 complex, is significantly reduced in *orc6* deletion fly larvae expressing GFP-Orc6-DmK23E mutant protein compared to the wild type GFP-DmOrc6. P - insoluble pellet after salt extraction, T- total ovaries extract, Pnut- loading control.

### Supplementary figure 2

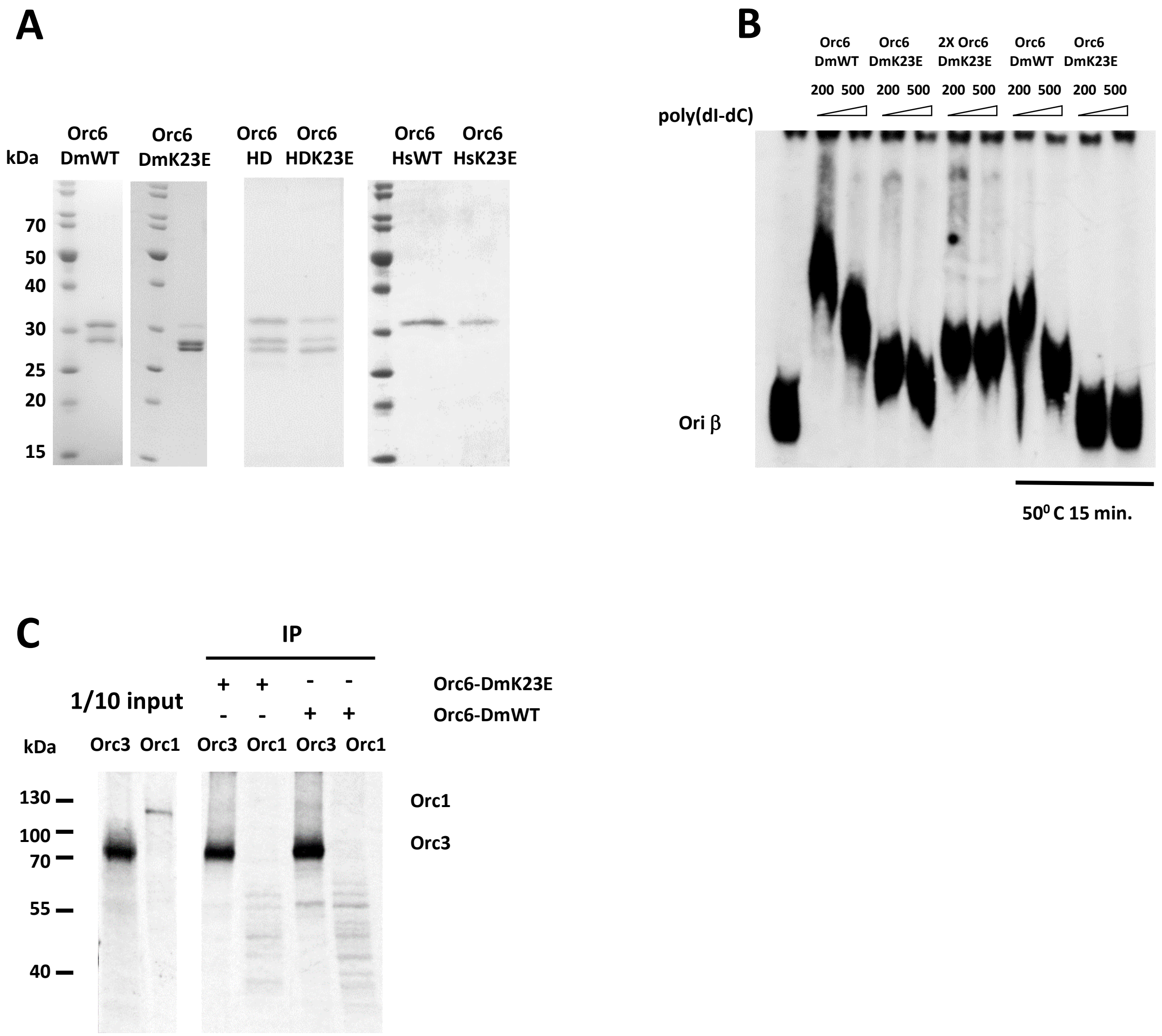


**(A)** Coomassie blue stained SDS gel of purified proteins used for electrophoretic mobility shift assays in Fig.6. **(B)** Electrophoretic mobility shift assay of *Drosophila* Orc6 wild type and K23E mutant, 2X Orc6-DmK23E indicates two fold increase in the amount of Orc6 protein in the reaction. '50°C 15 min.' indicates heat treatment of the protein for 15 minutes before adding to the reaction. **(C)** IP from *in vitro* transcription- translation reactions of Orc3 or Orc1 with anti-Orc6 antibodies. Purified recombinant *Drosophila* Orc6 wild type or K23E mutant proteins were used.

### Supplementary figure 3

**Orc6-DmWT** **Orc6-DmY225S**

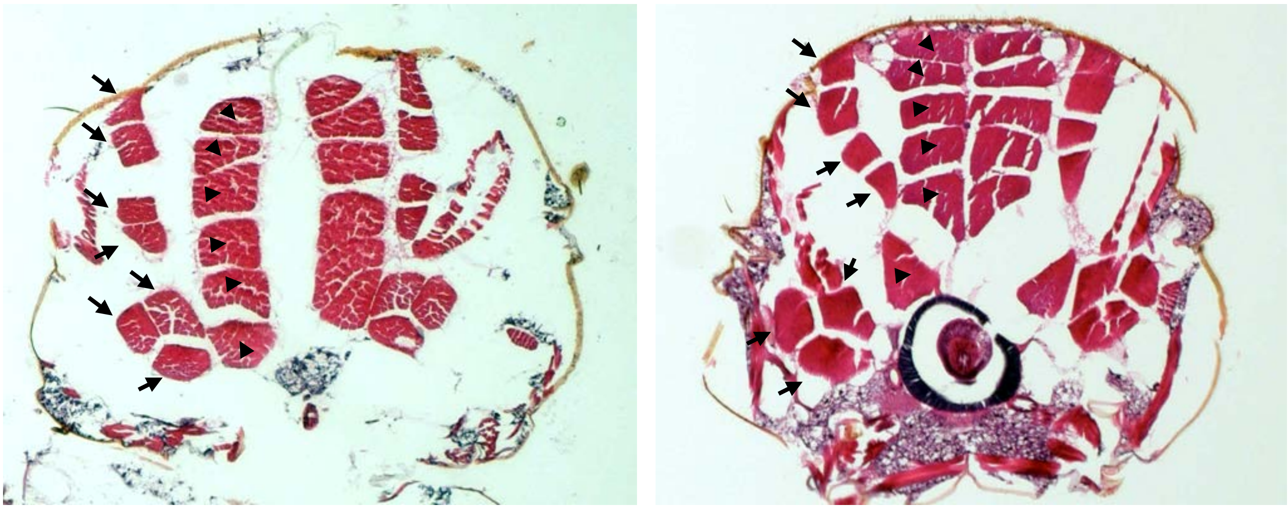

**Histological analysis of indirect flight muscles.** Adult flies were fixed and processed according to paraffin-embedded protocol (Kucherenko et al. 2010). Transversal paraffin sections of *Drosophila* thoraxes were selected to visualize indirect flight muscles (IFMs). IFMs are constituted of six dorsal longitudinal muscles (marked by arrowheads) and seven dorsoventral muscles (pointed by arrows) (Bernard et al. 2003). No muscle atrophy was detected in Meier-Gorlin flightless mutant compared to wild type. Orc6-DmWT – *orc6* deletion rescued with *Drosophila* wild type *orc6* transgene. Orc6-DmY225S – *orc6* deletion rescued with *Drosophila* Meier-Gorlin *orc6-DmY225S* transgene.

### Supplementary Table 1

#### Gene ontology annotation for Y225S mutant

| GO terms: DOWNREGULATED | GO terms: UPREGULATED |
|---|---|
| neurogenesis | proteolysis |
| mitotic nuclear division | transmembrane transport |
| DNA replication | oxidation-reduction process |
| DNA repair | innate immune response |
| transcription, DNA-templated | response to bacterium |
| oogenesis | circadian rhythm |
| DNA replication initiation | hexose transmembrane transport |
| cellular response to starvation | glucose import |
| mitotic spindle organization | carbohydrate metabolic process |
| rRNA processing | response to fungus |
| protein import into nucleus | transport |
| mitotic cell cycle | peptide catabolic process |

#### Genes that are downregulated in both Y225S and K23E mutants more then 2 times

| Drosophila Entrez Gene ID | Human Entrez Gene ID(s) | Symbol | Description | Maximum expression | Y225S vs wild-type | K23E vs wild-type |
|---|---|---|---|---|---|---|
| 251711 | | CG31517 | no data | Ovary | -4.5 | -1.7 |
| 41254 | 51548 | Sirt6 | Histone deacetylase | Ovary | -4.2 | -1.5 |
| 40217 | 23028 | Su(var)3-3 | Histone demethylase | Ovary | -3.8 | -2.0 |
| 318225 | | CG32816 | no data | Larval trachea | -3.0 | -2.0 |
| 42183 | 84154 | Non3 | rRNA binding activity | Ovary | -2.8 | -1.6 |
| 317935 | | CG32243 | no data | Ovary | -2.7 | -1.9 |
| 41994 | | CG14882 | Methionine synthase reductase | Ovary | -2.7 | -1.6 |
| 2768669 | 55425 | CG33332 | no data | Ovary | -2.4 | -1.5 |
| 42056 | 24140 | CG5220 | tRNA methyltransferase | Ovary | -2.3 | -1.9 |

#### Genes that are upregulated in both Y225S and K23E mutants more then 2 times

| Drosophila Entrez Gene ID | Human Entrez Gene ID(s) | Symbol | Description | Maximum expression | Y225S vs wild-type | K23E vs wild-type |
|---|---|---|---|---|---|---|
| 43248 | | CG17192 | Phospholipase A(1) | Digestive system | 5.4 | 11.7 |
| 43264 | | CG6074 | Carbonic anhydrase | Adult salivary gland | 3.8 | 3.7 |
| 43543 | | Jon99Ciii | Serine protease | Digestive system | 3.5 | 2.2 |
| 38240 | | Cpr62Bb | Cuticular protein | Adult head | 3.4 | 2.3 |
| 35498 | | TpnC4 | Troponin C isoform 4 | Adult carcass | 2.9 | 2.1 |
| 33707 | | Jon25Bii | Serine protease | Digestive system | 2.9 | 2.6 |
| 33708 | | Jon25Bi | Serine protease | Digestive system | 2.9 | 2.3 |
| 33706 | | Jon25Biii | Serine protease | Digestive system | 2.8 | 2.2 |
| 42451 | | CG4000 | no data | Adult head | 2.7 | 2.4 |
| 35859 | 57115 | PGRP-SC1a | Peptidoglycan recognition protein | Digestive system | 2.6 | 5.4 |
| 38328 | | CG14949 | no data | Digestive system | 2.6 | 2.8 |
| 34055 | | CG7203 | Cuticular protein | Adult eye | 2.2 | 2.7 |
| 38946 | | CG17352 | no data | Adult eye | 2.1 | 2.0 |
| 318098 | 91445 | CG32581 | Ubiquitin protein ligase | Adult head | 2.1 | 48.1 |
